# Ectopic methylation of a single persistently-unmethylated CpG in the promoter of the vitellogenin gene abolishes its inducibility by estrogen through attenuation of USF binding

**DOI:** 10.1101/768697

**Authors:** Lia Kallenberger, Rachel Erb, Lucie Kralickova, Andrea Patrignani, Esther Stöckli, Josef Jiricny

## Abstract

The enhancer/promoter of the vitellogenin II (*VT*G) gene has been extensively studied as a model system of vertebrate transcriptional control. While deletion mutagenesis and *in vivo* footprinting identified the transcription factor (TF) binding sites governing its tissue specificity, DNase hypersensitivity- and DNA methylation studies revealed the epigenetic changes accompanying its hormone-dependent activation. Moreover, upon induction with estrogen (E_2_), the region flanking the estrogen-responsive element (ERE) was reported to undergo active DNA demethylation. We now show that although the *VTG* ERE is methylated in embryonic chicken liver and in LMH/2A hepatocytes, its induction by E_2_ was not accompanied by extensive demethylation. In contrast, E_2_ failed to activate a *VTG* enhancer/promoter-controlled luciferase reporter gene methylated by *Sss*I. Surprisingly, this inducibility difference could be traced not to the ERE, but rather to a single CpG in an E-box (CACGTG) sequence upstream of the *VTG* TATA box, which is unmethylated *in vivo*, but methylated by *Sss*I. We demonstrate that this E-box binds the upstream stimulating factor USF1/2. Selective methylation of the CpG within this binding site with an E-box-specific DNA methyltranferase *Eco*72IM was sufficient to attenuate USF1/2 binding *in vitro* and abolish the hormone-induced transcription of the *VTG* gene in the reporter system.

## INTRODUCTION

In vertebrates, the DNA methyltransferases DNMT1, DNMT3a and DNMT3b convert around 90% of cytosines in the CpG sequence context to 5-methylcytosines (Eckhardt et al., 2006; Gruenbaum et al., 1981; Rakyan et al., 2004). DNMT3a/b are believed to be the major *de novo* methyltransferases that modify unmethylated DNA, while DNMT1, often termed “maintenance methylase”, is believed to be the enzyme that copies the methylation pattern of the template strand onto the newly-synthesised strand following DNA replication or repair (Bacolla et al., 1999; Bestor, 1992; Hsieh, 1999; Okano et al., 1998; Pradhan et al., 1999). All three enzymes are essential for survival, as demonstrated by the fact that DNMT knock-out mice show early lethality (Li et al., 1992; Okano et al., 1999).

DNA methylation is largely erased during fertilisation, but the DNMTs lay down a new methylation pattern during early embryogenesis that will control the subsequent stages of development and differentiation. In general, gene bodies become densely-methylated, while gene regulatory sequences are methylated sparsely and in a highly-divergent manner. For example, many housekeeping genes are flanked by the so-called CpG islands. Although these regions are CpG-rich, they are generally unmethylated and the genes they control are constitutively-active (Bird et al., 1985; Cooper et al., 1983; Gardiner-Garden and Frommer, 1987). In contrast, CpG islands associated with imprinted genes or retroviral sequences are methylated, as are genes on the inactive X chromosome (Liu et al., 1994; Walsh et al., 1998; Woodcock et al., 1997) and some become methylated during development (De Smet et al., 1996; De Smet et al., 1999), which leads to transcriptional silencing. Once established, DNA methylation patterns remain largely stable and unprogrammed changes such as the aberrant methylation of CpG islands are often linked to aging or tumorigenesis (Baylin et al., 1986; Gama-Sosa et al., 1983; Goelz et al., 1985; Noreen et al., 2014; Toyota et al., 1999). While the latter phenomena have been extensively studied, less attention has been paid to the dynamic changes of DNA methylation taking place outside of CpG islands (Eden and Cedar, 1994). These changes are often triggered by exogenous stimuli in a highly tissue-specific manner and are directly involved in the regulation of gene expression (Amenya et al., 2016; Kangaspeska et al., 2008; Metivier et al., 2008; Thomassin et al., 2001; Toker et al., 2013) by altering the binding affinity of TFs such as c-Myc/Myn (Prendergast and Ziff, 1991), E2F (Campanero et al., 2000), AP2 (Comb and Goodman, 1990), NF-κB (Kirillov et al., 1996) or USF1/2 (Fujii et al., 2006) for their cognate sequences.

One well-studied example of an inducible tissue-specific gene that is also regulated by DNA methylation is vitellogenin II (*VTG*). The gene encodes a precursor of egg yolk protein and is present in all oviparous species. It is expressed exclusively in the female liver, but can be induced in males by estrogen (Saluz et al., 1986). This property brought it into recent limelight, because its expression in males can be used as a measure of estrogenic endocrine disruptive chemicals (EDCs) in the environment (Diamanti-Kandarakis et al., 2009). As in other species, the chicken *VTG* gene is expressed in the liver of mature hens, but not roosters. This difference was explained by the silencing of the *VTG* gene by sex-specific DNA methylation, because its transcriptional activation in rooster liver by a single β-estradiol (E2) injection was accompanied by demethylation of a *Hpa*II site within the estrogen response element (ERE) (Wilks et al., 1984; Wilks et al., 1982) and the appearance of DNaseI hypersensitive sites in the enhancer and promoter (Burch and Weintraub, 1983). Subsequent Church & Gilbert sequencing of the genomic DNA showed that the transcription was activated already after 6 hours and that this event coincided with the demethylation of four CpGs (**a**-**d**) in the non-transcribed strand flanking the ERE (Fig. 1A). Because loss of methylation through replication (the so-called “passive demethylation”) could be excluded, this phenomenon was hailed as the first example of active demethylation (Saluz et al., 1986).

**Fig 1.**
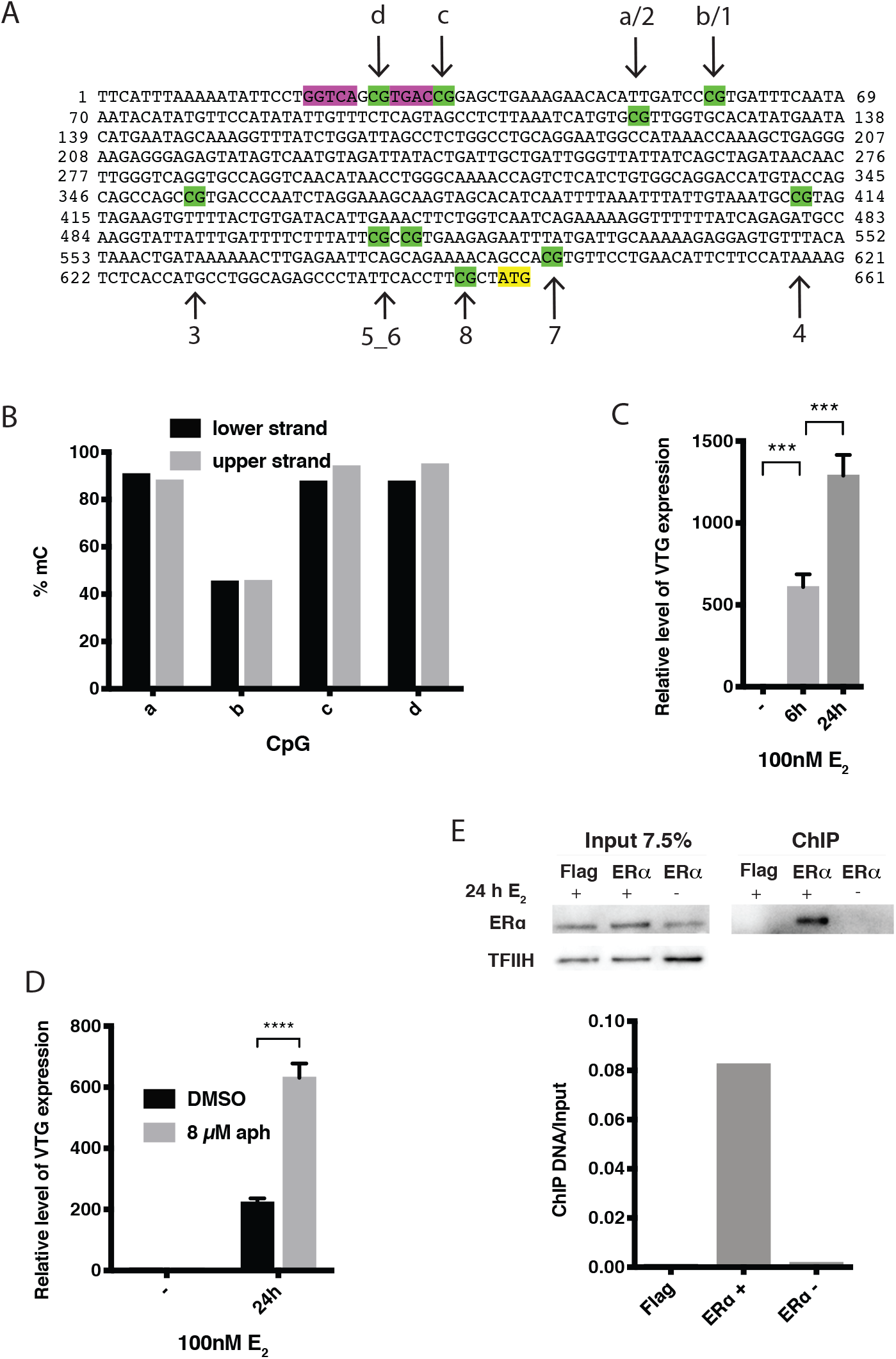
Sequence of the *VTG* enhancer/promoter, its methylation and inducibility *in vivo.* **A**, Sequence of the enhancer/promoter region of the chicken *VTG* gene. The ERE binding site (violet), the CpGs (green) and the translation start site (yellow) are highlighted. The four CpGs (**a**-**d**) analyzed in Saluz *et al.* (Saluz et al., 1986) are indicated, as well as the additional six CpGs (3-8) in the enhancer/promoter region. **B**, Bisulphite sequencing of CpGs **a**-**d** in LMH/2A cells. **C**,**D** *VTG* mRNA levels measured by RT-qPCR after 6 and 24 hours of 100 nM E_2_ treatment (**C**) and upon additional treatment with 8 µM aphidicolin (aph) or DMSO for 24h (**D**). Data are represented as mean ± SD. Significance was assessed using Sidak’s multiple comparisons test. *P≤0.05, **P≤0.01, ***P≤0.001, ****P≤0.0001. **E**, Upper panel: Western Blot of chromatin extracts immunoprecipitated with Flag– or ERα antibodies (- and + β-estradiol treatment). Lower panel: Ratio of RT-qPCR signal of immunoprecipitated versus input DNA.

We set out to study the above phenomenon in greater detail, because we wanted to learn whether the demethylation was an obligate step in the activation of *VTG* expression and, if so, whether it involved the recently-discovered machinery of active DNA demethylation that makes use of the ten-eleven-translocation methylcytosine dioxygenase (TET) enzymes (Hassan et al., 2017; He et al., 2011; Tahiliani et al., 2009) and TDG (Neddermann and Jiricny, 1993; Wiebauer and Jiricny, 1989). We made use of chicken embryos (Burch and Weintraub, 1983), or of a chicken hepatoma LMH/2A cell line stably-expressing the estrogen receptor (ERα), (Binder et al., 1990; Philipsen et al., 1988; Seal et al., 1991; Sensel et al., 1994), both of which had been used to study *VTG* expression in the past. We also made use of a reporter plasmid that was devoid of CpGs and in which the expression of the luciferase gene was under the sole control of the *VTG* enhancer/promoter. We now show that estrogen-dependent *VTG* activation in these experimental systems was not accompanied by significant demethylation. More importantly, we show that the ability of the gene to reactivate from methylation-dependent silencing is controlled by an unmethylated E-box element distal to the ERE sequences. Methylation of this E-box abolished the ability of the silenced gene to be reactivated by estrogen. We further show that the E-box binds the upstream-stimulating factor USF1/2.

## RESULTS

### The *VTG* enhancer/promoter is methylated and the gene is silenced, but exposure to β-D-estradiol induces transcription independently of DNA replication

We first wanted to reproduce the phenomenon described by Saluz *et al.* (Saluz et al., 1986). However, due to restrictions on animal experimentation, we had to search for alternative systems. We were also interested in identifying an experimental set-up that would be amenable to manipulation. To this end, we decided to test whether the *VTG* gene is silenced and inducible in the liver of 10 day-old chicken embryos. We made small windows in freshly-fertilised eggs, sealed them with a microscope coverslip and hot wax (Baeriswyl and Stoeckli, 2006) and incubated them at 39°C for 9 days. Eggs containing live embryos were then treated with an ethanol solution of E_2_ or with ethanol alone. 24 hours later, the embryos were sacrificed, the livers were excised, immersed in RNA*later* and nucleic acids and proteins were immediately isolated. In parallel, we treated chicken LMH/2A cells, a Leghorn rooster hepatocellular carcinoma cell line stably expressing estrogen receptor alpha (ERα) (Sensel et al., 1994) in a similar manner, but nucleic acids and proteins were isolated after 6 and 24 hours.

Bisulphite sequencing of DNA isolated from mock-treated liver and LMH/2A cells revealed that CpGs a/2, c and d in the *VTG* enhancer (Fig. 1A) were ∼90% methylated in both strands, whereas CpG b/1 appeared to be undermethylated in both LMH/2A cells (Fig. 1B) and embryonic liver (Fig. S1A). This is in line with the results of Saluz *et al*., where CpGs a/2, c and d in rooster liver DNA were reported to be fully-methylated, whereas CpG b/1 was hemimethylated (Saluz et al., 1986).

Using RT-qPCR, we could show that the *VTG* gene was transcribed neither in LMH/2A cells nor in embryonic liver prior to exposure to estrogen, but that it was induced upon a 24 hours E_2_ treatment of chicken embryos (Fig. S1B) and even more efficiently in LMH/2A cells, where substantial transcription was detected already 6 hours after the addition of 100 nM E_2_ (Fig. 1C). Its expression increased further after 24 hours and the increase continued up to 72 hours (Fig. S1C), possibly due to a positive feedback leading to increased transcription of ERα (Fig. S1D). This activation occurred in a replication-independent manner, as inhibition of DNA replication by the addition of the B-family polymerase inhibitor aphidicolin did not impede transcription, but rather enhanced it (Fig. 1D). A possible explanation for this observation is that interference of replication with the transcription process is inhibited by the addition of aphidicolin. That DNA synthesis was indeed inhibited was confirmed by a lack of 5-ethynyl-2’-deoxyuridine (EdU) incorporation into nuclear DNA during the course of the experiment (Fig. S1E).

### Estrogen-dependent induction of *VTG* transcription in our system was not accompanied by demethylation of the enhancer/promoter region

We next set out to investigate whether the estrogen-dependent transcriptional activation of the *VTG* gene was accompanied by demethylation as reported (Saluz et al., 1986). We therefore performed PCR on the bisulphite-converted DNA to distinguish cytosines (C) from 5-methylcytosines (mC). Because active demethylation is believed to involve the dioxygenases TET1-3, we also wanted to detect the intermediate of the oxidation, 5-hydroxymethylcytosine (hmC). To distinguish between mC and hmC, the bisulphite conversion was preceded by an oxidation step using KRuO_4_, which selectively oxidizes hmC to fC, resulting in its conversion to uracil following bisulphite treatment (Booth et al., 2012). The PCR products were then sequenced using PacBio, which provides information on single molecules. HmC levels were assessed as the difference in the amount of converted cytosines between bisulphite- and oxidized-bisulphite converted DNA. Using this approach, we could show that the *VTG* promoter/enhancer was not demethylated during the course of the induction, but that a small percentage of mCs were oxidized to hmCs, indicating a possible involvement of the TET enzyme(s) (Fig. S1F).

### ERα binds to the *VTG* estrogen response element upon treatment with β-estradiol *in vivo*

As mentioned in the Introduction, DNA methylation can attenuate transcription by interfering with the binding of TFs to their respective recognition sequences. Because we failed to detect demethylation in our system, ERα, the key activator of the *VTG* gene, would have had to bind to the methylated estrogen response element (ERE) in the enhancer upon E_2_ treatment. In order to test this hypothesis, we performed a chromatin immunoprecipitation experiment using an ERα antibody or, as a control, a flag antibody. We could retrieve ERα in the chromatin fraction only with the ERα antibody and upon E_2_ treatment (Fig. 1E, upper panel). Moreover, we could confirm its binding to the *VTG* ERE using RT-qPCR on the recovered DNA (Fig. 1E, lower panel). Only very little signal was apparent without E_2_ treatment, confirming the widely-accepted model that estrogen receptors bind DNA *in vivo* only in response to hormone treatment (Klinge, 2001). Moreover, this showed that the receptor is able to bind to its recognition sequence also in methylated chromatin.

### Binding of ERα to the ERE is insensitive to different cytosine modifications and hormone treatment

To confirm that the binding of ERα to its cognate recognition sequence was indeed unaffected by methylation, we carried out a series of electrophoretic mobility shift assays (EMSAs), using synthetic oligonucleotides containing the *VTG* ERE (WT) or a variant with a single base pair deletion in the three-nucleotide spacer between the palindromic repeats (ΔG, Fig. 2A) that has been reported to abolish ERα binding *in vitro* (Klinge, 2001). Recombinant ERα was able to bind the ERE sequence with similar affinity in the presence or absence of E_2_, the only notable difference being the slightly-increased mobility of the shifted band in the presence of the hormone (Fig. 2A, lanes 2,3 and Fig S2A,B). The specificity of the protein/DNA complex was confirmed by a competition assay; addition of a 100-fold excess of the unlabelled ERE oligo duplex significantly diminished the intensity of the shifted band (lane 4, spec), while a scrambled sequence failed to do so (lane 5, unspec). In addition, addition of an ERα antibody to the reaction substantially retarded (supershifted) the mobility of the specific band (lane 7) as compared to the addition of the same amount of BSA (lane 6). In the EMSA assays, the affinity of ERα for a substrate symmetrically-methylated at the two CpGs within the ERE (CpGs c and d, mC/mC) was similar to that seen with the unmethylated substrate C/C (Fig. 2B) and the same was true for hemi- and fully-hydroxymethylated substrates, as well as substrates containing formyl- or carboxycytosine (Fig S2C,D).

**Fig 2.**
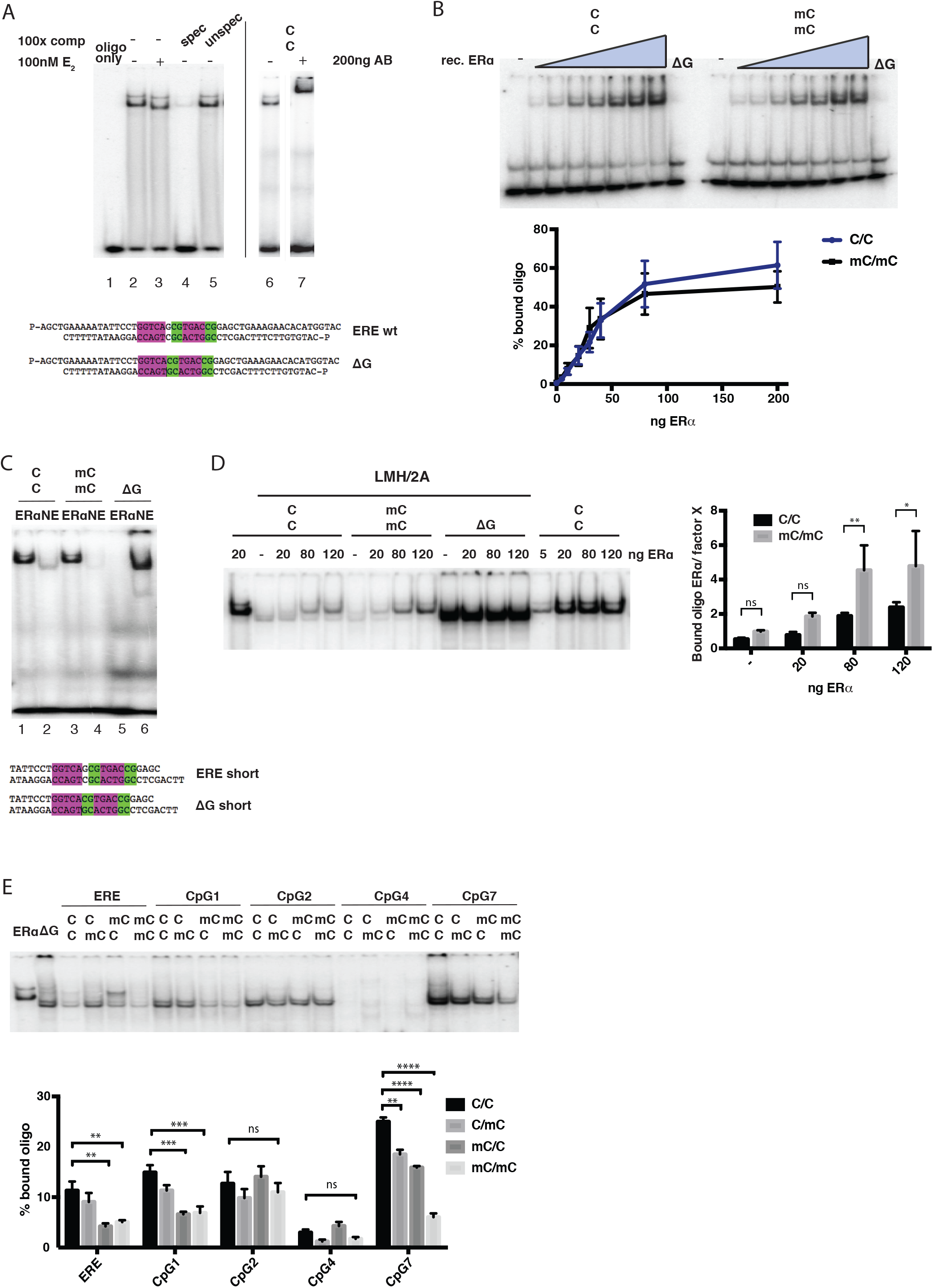
Electrophoretic mobility shift analysis of ERE-binding proteins. **A**, EMSA with 60 ng recombinant ERα and unmethylated ERE wt oligo (C/C), with (+, lane 3) or without (-, lanes 1, 2, 4-7) E_2_ or competitor. The specific competitor (spec, lane 4) was the unlabelled ERE wt duplex C/C, the unspecific competitor (unspec, lane 5) was an unrelated duplex of similar length. An antibody against ERα (MA5-13065, lane 7) or BSA (lane 6) were used in the supershift experiment with the unmethylated oligo C/C. The oligonucleotides used for the EMSAs are depicted below the autoradiograph. Purple, consesus ERE; green, CpGs. The ΔG duplex contains a single base pair deletion (white) in the spacer region between the two dyad-symmetry elements of the ERE. **B**, Upper panel: representative EMSAs with increasing concentrations of recombinant ERα (0, 5, 10, 20, 30, 40, 80 and 200 ng) and unmethylated (C/C, left panel) or methylated (mC/mC, right panel) ERE wt or ΔG oligo with 40 ng of ERα. (In the mC/mC duplex both CpGs were symmetrically-methylated.) Lower panel: Quantification of percentage bound oligo in three independent experiments. Data are represented as mean ± SD. **C**, EMSA comparing the binding of recombinant ERα protein or LMH/2A NEs to shorter duplexes (ERE short shown below the autoradiograph) unmethylated (C/C) and methylated (mC/mC) or a shorter ΔG oligo (ΔG short). **D**, Left panel: EMSA with the indicated duplexes and LMH/2A NE supplemented with the indicated amounts of recombinant ERα. Right panel: Quantification of the ratio of oligo bound by ERα or factor X in three independent experiments. Data are represented as mean ± SD. Significance was assessed using the Holm-Sidak test. **E**, Top panel: EMSA with LMH/2A nuclear extracts and five duplexes containing the indicated CpGs from the *VTG* enhancer/promoter (Fig. 1A), bearing different combinations of cytosines and methylcytosines. Bottom panel: Quantification of percentage bound oligo in three independent experiments. Data are represented as mean ± SD. Significance was assessed using the Tukey test for multiple comparisons. The panels show autoradiographs of non-denaturing 6% polyacrylamide gels eluted with TAE buffer. *P≤0.05, **P≤0.01, ***P≤0.001, ****P≤0.0001.

### The ERE is bound by a factor other than ERα in nuclear extracts of LMH/2A cells

We were interested to learn whether the oligonucleotide substrates were bound with similar selectivity and affinity also by ERα present in nuclear extracts of LMH/2A cells. In order to limit non-specific binding, we used shorter substrates than in the previous experiments. As seen in Fig. 2C, the ERE substrates were efficiently bound by recombinant ERα irrespective of methylation (lanes 1, 3), whereas the ΔG oligonucleotide duplex failed to bind the receptor (lane 5). Unexpectedly, the mobility of the shifted band generated by the unmethylated oligo (C/C) upon incubation with LMH/2A nuclear extracts (NE) was faster than that seen with ERα (lane 2) and the factor that bound the unmethylated oligo bound only weakly to the methylated (mC/mC) one (lane 4). In contrast, the ΔG substrate was bound with very high affinity (lane 6). In order to ensure that the factor binding the oligo substrates in the extracts was distinct from ERα, we titrated increasing amounts of the recombinant protein into the NE. As shown in Fig. 2D (whole gel is shown in Fig S2E), ERα outcompeted the nuclear factor (referred to as Factor X) in reactions containing the C/C substrate and even more efficiently in reactions containing the mC/mC oligo, but not the ΔG substrate.

In order to gain information regarding the substrate preference of Factor X, we carried out an EMSA assay using eight oligonucleotide substrates carrying sequences flanking the ten CpGs in the *VTG* enhancer/promoter (Fig. 1A; the two CpGs in the ERE were in a single duplex, as were CpGs 5 and 6, because of their proximity). Because the sequence was reported by Saluz *et al.* (Saluz et al., 1986) to be demethylated in a strand-specific manner, we also included hemimethylated substrates. Surprisingly, in addition to oligos ERE and ΔG, Factor X bound also to oligos containing CpG b/1, CpG a/2 and CpG7, but not CpG3, CpG4, CpG5-6 and CpG8 (Fig. 2E and Fig. S2F-H). Moreover, its affinity for hemimethylated substrates varied, which suggested that its sequence- and methylation specificity was unusually relaxed.

To characterize Factor X further, we carried out the EMSA assays with oligos containing one BrdU residue on each strand (Fig. 3A, Table S2). Upon incubation with the NE, half of the mixture was used for the EMSA experiment, while the other half was UV-crosslinked and the proteins were then resolved by SDS-PAGE. During the cross-linking, the radiolabelled oligo becomes covalently attached to the protein that binds it and the protein size can thus be estimated from the position of the radioactive band on the SDS-PAGE, once the molecular weight of the oligo is subtracted. As shown in Fig. 3B, the recombinant ERα-oligo complex migrated at the expected size of ∼80 kDa (lane 5) and a band of similar size was seen in LMH/2A NE cross-linked to the C/C, mC/C or mC/mC substrates (lanes 1-4). In addition, a second, prominent band migrated at around 50 kDa, irrespective of substrate. After subtracting the molecular weight of the single-stranded oligo (∼15 kDa), the size of Factor X was predicted to be around 35-40 kDa. When the recombinant receptor was titrated into the reaction with NE, we saw a weak, but reproducible, competition with the smaller protein (Fig. 3C). The ∼80 kDa band could be outcompeted with a specific (lane 8, spec), but not with an unspecific (lane 7, unspec) competitor, further confirming that it was a complex of the oligo with ERα. The addition of E_2_ did not alter the binding affinities or ratios of the different proteins to the substrates (Fig. S3A). The migration of the ERα band was also unchanged in the presence of the hormone, confirming that the mobility shift seen in the EMSA assay represented a conformational change of the receptor, which is eliminated upon denaturation.

**Fig 3.**
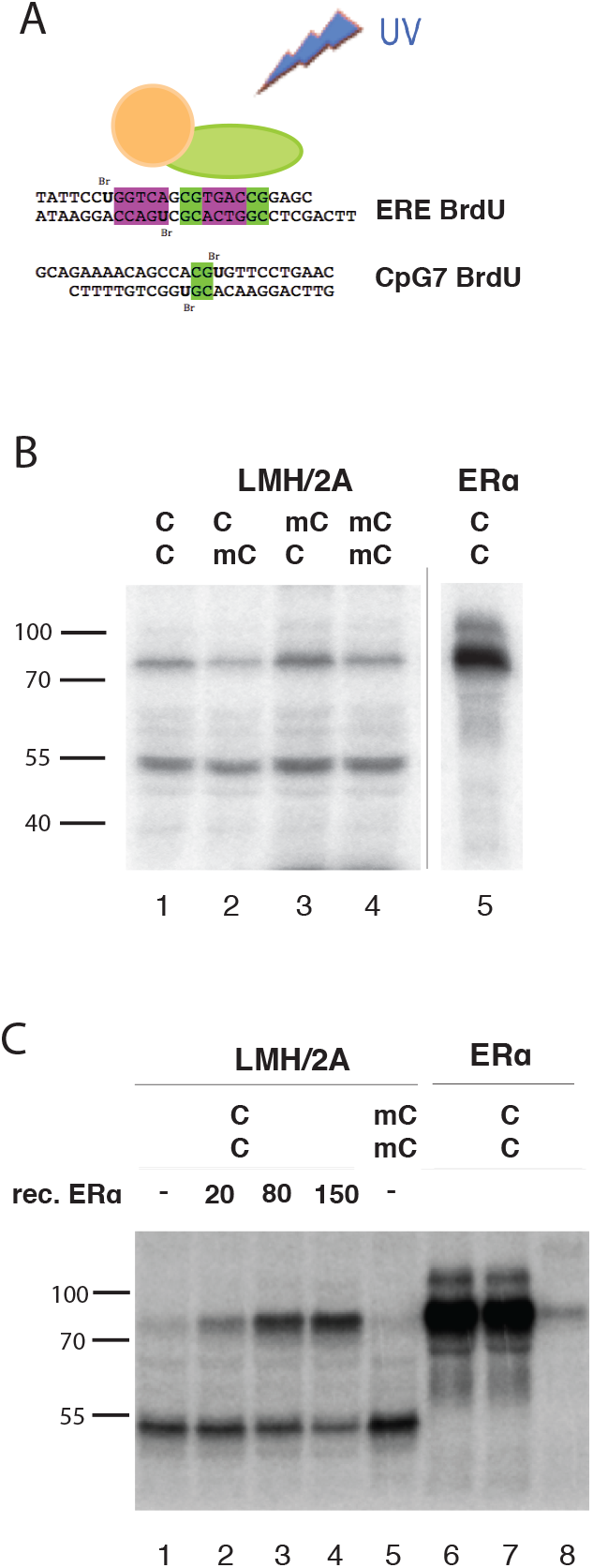
Identification of Factor X by UV cross-linking. **A**, Experimental set-up showing the position of the BrdU residues in the indicated duplex substrates. The binding reactions contained either 60 µg of LMH/2A NE or recombinant ERα. **B,** Proteins cross-linked to the oligo substrates shown in A (unmethylated or methylated as indicated). **C**, As in B, but the cross-linking reactions were supplemented with the indicated amounts of recombinant ERα or competitor oligo. The cross-linked complexes were resolved by 10% SDS-PAGE and visualised by autoradiography.

### Factor X is the E-box-binding heterodimer of upstream stimulating factors USF1/2

We wanted to learn whether Factor X was a chicken-specific protein, or whether it was present also in man. We therefore studied the binding properties of the eight oligonucleotide duplexes containing the ten CpGs in the *VTG* enhancer/promoter (see above) in EMSA experiments using NEs of the ERα-positive breast cancer cell line MCF7 and the ERα-negative cervical carcinoma cell line HeLa. As shown in Fig. S3B,C, the electrophoretic mobilities of the protein/DNA complexes formed in HeLa (ERE, ΔG, CpG4 and 7) and MCF7 (CpG4, 5-6, 7 and 8) extracts with the indicated oligos were similar to those seen in LMH/2A extracts. We therefore assigned them to Factor X. A distinct mobility shift was seen with oligo CpG3 (Fig. S2G), but the CpG4, 5/6 and 8 substrates failed to form protein/DNA complexes (Fig. 2E, Fig. S2H, Fig S3B,C). Interestingly, Factor X binding to the different oligos displayed distinct methylation sensitivities; thus, while its binding to CpG2 was unaffected by methylation, the binding to oligos ERE, CpG1 and in particular CpG7 was substantially attenuated by methylation (Fig. 2 E). The similarity of the proteins binding the oligo substrates ERE and CpG7 was further confirmed in UV-crosslinking experiments (Fig. S3D), thus implying that Factor X is a generic DNA binding protein with an ambiguous sequence- and methylation specificity.

In an attempt to identify Factor X, we subjected nuclear extracts of HeLa cells to affinity chromatography. We first generated the DNA substrates by ligating the ERE (C/C or mC/mC), CpG7 (C/C or mC/mC) or ΔG oligonucleotide duplexes end-to-end so as to obtain molecules of 100-300 base pairs in length. The sticky ends were then filled-in with dATP and Bio-dUTP and the tagged molecules were allowed to attach to streptavidin Dynabeads. We then preincubated HeLa NEs with non-specific competitor poly (dI-dC) and allowed them to incubate with the beads as described in Material and Methods. Following extensive washing, the bound proteins were eluted with high salt and the tryptic digests were analysed by mass spectrometry (Fig. 4 A). We searched for peptide sequences present preferentially in the CpG7 and ΔG but not in the methylated CpG7 elutions and originating from proteins with molecular size between 25 and 50 kDa. Beads-only and CpG8 were used as negative controls. The peptides that fulfilled these exclusion criteria to the greatest extent were the TFs DEC1, MAX, MLX, Myc, USF1 and USF2, firstly because they were identified with elevated frequencies (Table 1) and, second, because their preferred recognition sequences overlapped to a large extent with those of our affinity probes.

**Figure 4.**
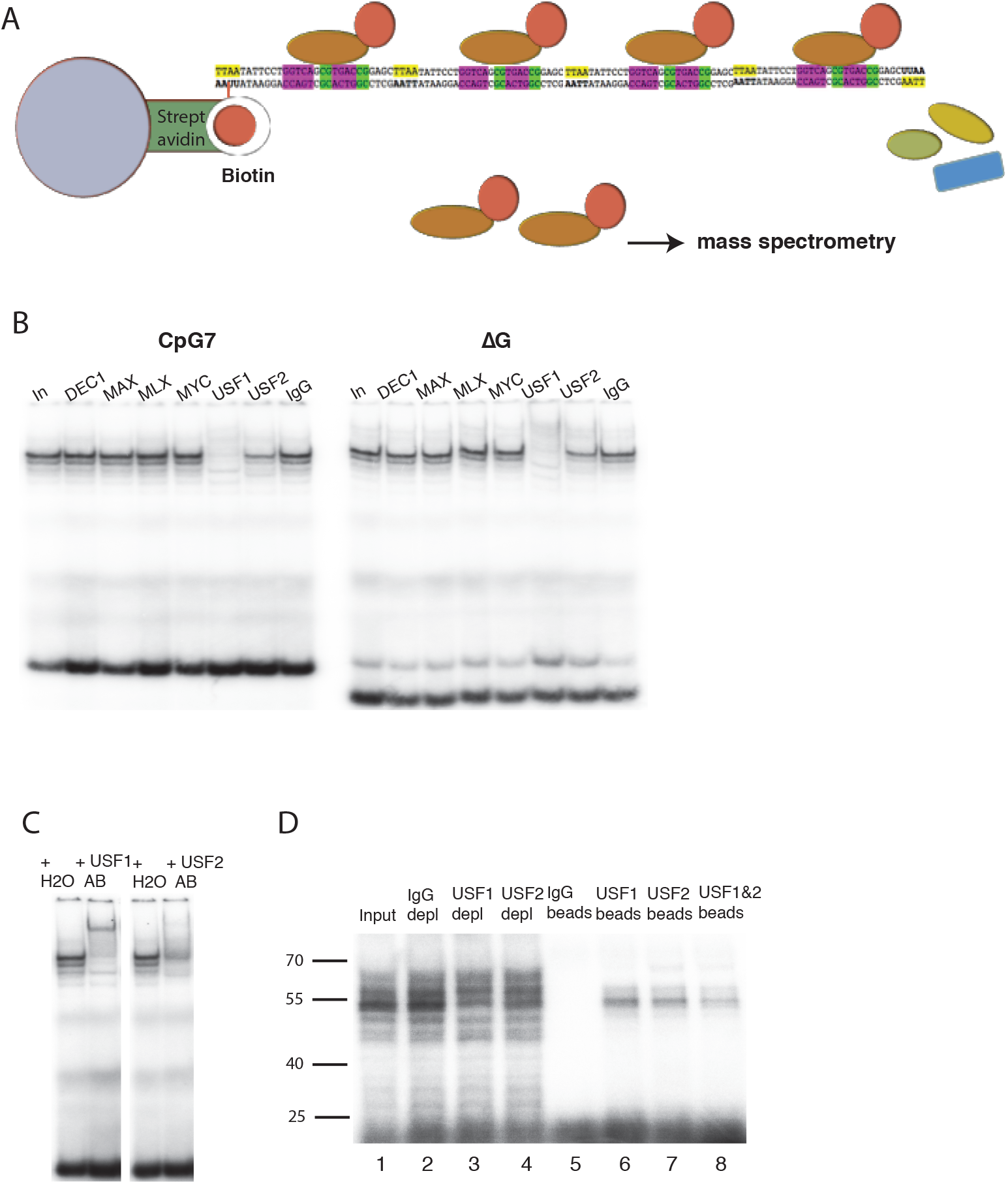
Identification of Factor X by affinity chromatography/MS. **A**, Experimental set-up of affinity pull-down and mass spectrometric analysis. The oligonucleotide duplexes were ligated end-to-end, tailed with bio-dUMP and bound to streptavidin Dynabeads. The beads were incubated with the extracts and the eluted proteins were analysed by MS as described in Materials and Methods. **B**, EMSA with oligo duplex CpG7 (left panel) or ΔG (right panel) and HeLa NE preincubated with the indicated antibodies. **C**, Supershift using oligo duplex CpG7 and antibodies specific for USF1 or USF2. **D,** UV cross-linking reactions. Depleted extracts (lanes 1-4) and proteins eluted from the Dynabeads (lanes 5-8) were bound to the unmethylated ERE oligo, UV-cross-linked, separated on SDS-PAGE and visualized by autoradiography.

**Table 1.**
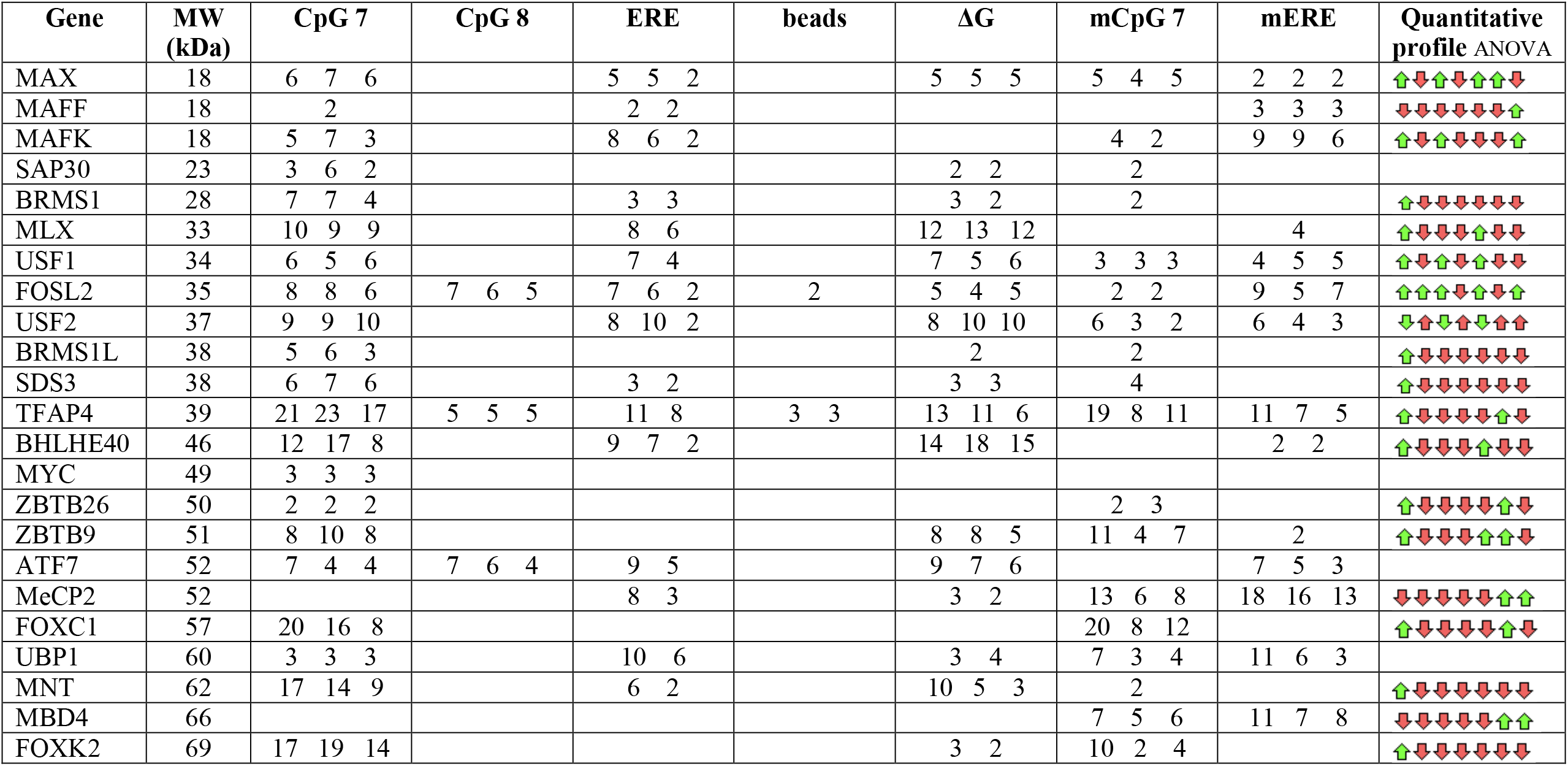
Peptides identified by LC-MS/MS in fractions eluted from affinity chromatography on the indicated oligonucleotide matrices. To subtract proteins that bound unspecifically to the matrix, beads only were used. As a second filter, we used an oligonucleotide containing the sequence around CpG8 that showed no binding of factor X in the EMSA experiments. Significance was assessed with an ANOVA multiple comparison t-test. Only hits identified with >95% confidence are shown. The false discovery rate (FDR) threshold was set to 1%.

We immunodepleted HeLa nuclear extracts with antibodies against these possible Factor X candidates and performed EMSA assays. The USF1-or USF2-immunodepleted extracts (Fig. S4A) largely lost the ability to bind the CpG7 and ΔG duplexes and the fraction of the shifted oligos in the USF2-depleted extracts was clearly reduced (Fig 4B). The same could be shown with the unmethylated- and methylated ERE substrates (Fig. S4B). For a direct comparison, Fig. S4C shows an EMSA experiment in which the four oligo substrates were incubated with the USF1-depleted and IgG-depleted (negative control) extracts. To confirm the depletion specificity, we performed EMSAs with HeLa NE that were pre-incubated with antibodies raised against the above proteins. As shown in Fig. 4C, the antibody against USF1 caused a supershift of the complex, while the antibody against USF2 attenuated the binding of Factor X to the two duplexes. To confirm the specificity of the depletion further, we carried out a UV-crosslinking experiment with the USF1-or USF2-depleted extracts, as well as with proteins eluted from the beads with high salt. The effect of the depletion was not very pronounced (Fig. 4D, lanes 1-4), probably because EMSA assays detect complexes with relatively long life times, whereas UV-cross-links even transient complexes. However, a band corresponding in size to Factor X was present in the eluates from the USF1 and USF2 beads, but not from the control, IgG beads (Fig. 4D). This evidence strongly suggests that Factor X is a heterodimer of USF1 and USF2, which is known to be methylation-sensitive (Chen et al., 2012; d’Adda di Fagagna et al., 1995; Fujii et al., 2006). However, our data provide strong evidence that this heterodimer is rather promiscuous in its sequence specificity, contrary to reports that describe its preference for the perfect E-box consensus CACGTG (d’Adda di Fagagna et al., 1995; Giacca et al., 1989) present e.g. in the CpG7 duplex.

### Transcriptional activity of the *VTG* enhancer/promoter is controlled by E-box methylation and USF binding

In the Saluz *et al*. study (Saluz et al., 1986), the silenced, methylated *VTG* gene was activated by estrogen and this event was accompanied by demethylation of CpGs **a**-**d** in the enhancer. These findings led to the assumption that the E_2_-activated estrogen receptor triggered a series of events that brought about a demethylation of the ERE and the downstream sequences, and that this demethylation was a prerequisite for transcriptional activation. However, in our *in vivo* system, induction of *VTG* transcription with E_2_ did not require demethylation (Fig. 1C and Fig S1F). We therefore had to consider the possibility that the gene was silent in rooster only because of the lack of the hormone and that its methylation was simply a mark of inactive chromatin. In order to elucidate this phenomenon, we generated a reporter vector based on the CpG-free pCpGL-basic plasmid, in which the luciferase gene is driven by the *VTG* enhancer/promoter. We first introduced into the *VTG* sequence unique *Kpn*I and *Hin*dIII sites on either side of the ERE and then ligated this enhancer/promoter into pCpGL-basic to generate VTG-CpGL (Fig S5A). Upon transfection of this reporter into LMH/2A cells, luciferase expression could be efficiently induced with E_2_ (Fig 5A, ERE). Convertion of the wild type ERE sequence to ΔG by site-directed mutagenesis resulted in similar levels of basal transcription, but substantially lower inducibility (Fig. 5A), caused by the significantly lower affinity of ERα for the ΔG ERE lacking the spacer deoxyguanosine (Fig. 2B,C). *In vitro* methylation of these two reporters with *Sss*I largely abolished transcriptional inducibility (Fig. 5A). When the *Kpn*I/*Hin*dIII fragment was replaced with the wild type sequence, unmethylated, methylated or hydroxymethylated at the two CpGs, estrogen inducibility was unchanged (Fig. 5B). This result extends the *in vitro* findings (Fig. 2B, Fig. S2C) showing that the binding of ERα is unaffected by methylation and shows that its transcriptional activity is also methylation-independent. (Hydroxymethylation was included, because we detected low levels of this modification in the *in vivo* activation experiments shown in Fig. S1F.)

**Figure 5.**
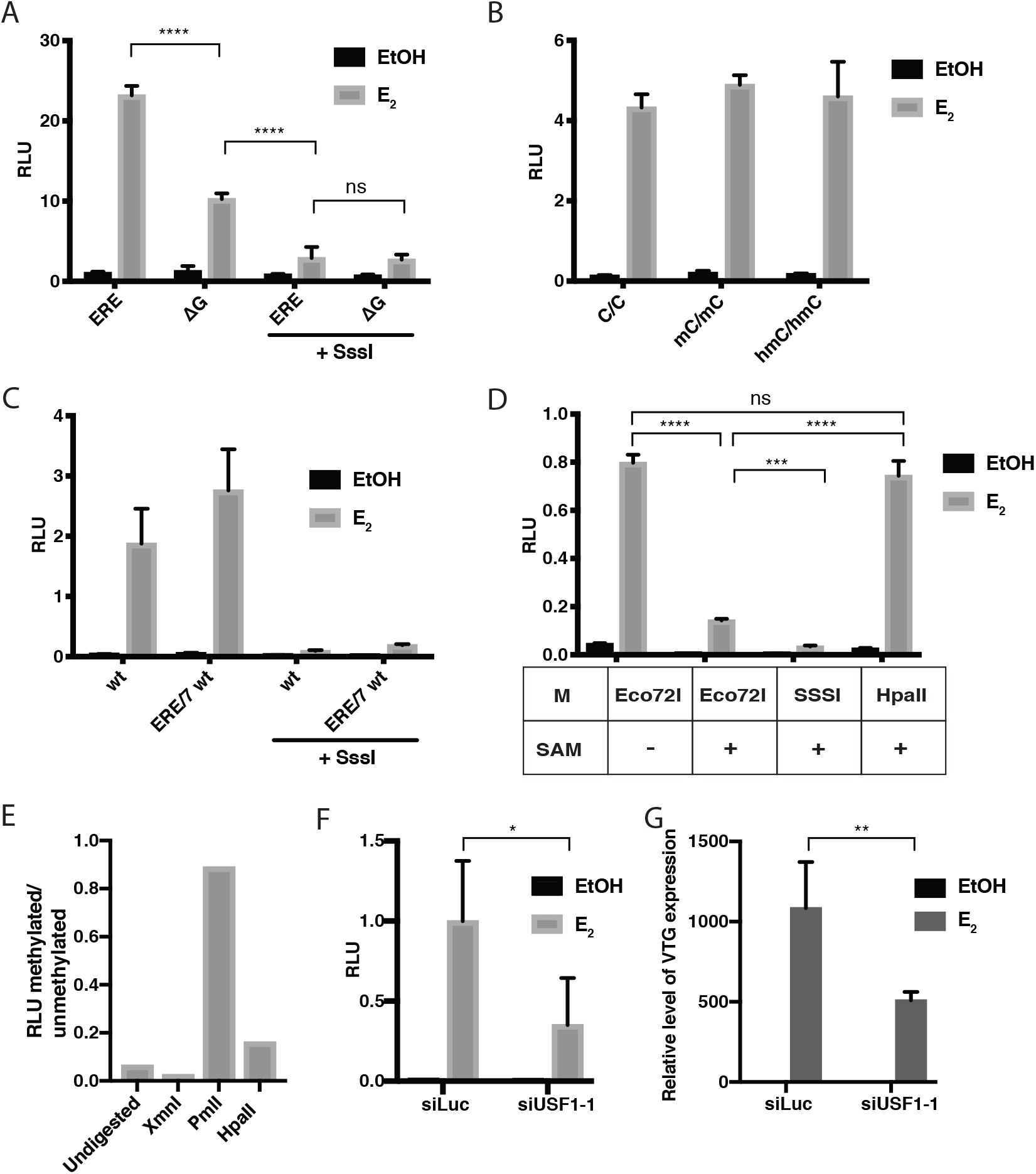
Estrogen inducibility of VTG-CpGL luciferase reporter vector transfected into LMH/2A cells. **A**, Luciferase expression before and after *in vitro* methylation of the ERE C/C (wt) or ΔG vectors with *Sss*I. **B**, Same as A, but the ERE sequence was replaced with oligonucleotides carrying the indicated cytosine modifications only in CpGs **d** and **c**. **C**, Same as A, but either with wt VTG-CpGL or the mutated reporter in which all CpGs except for the ERE and CpG7 were substituted for TpGs, +/-*Sss*I. **D**, Luciferase expression from the reporter before and after methylation with *Eco*72IM, *Hpa*II.M or *Sss*I. *Eco*72IM without S-adenosylmethionine (-SAM) was used as control, the other methylation reactions were performed in the presence of SAM (+SAM). Significance was assessed using the Tukey test for multiple comparisons. *P≤0.05, **P≤0.01, ***P≤0.001, ****P≤0.0001. **E**, Luciferase assay using VTG-CpGL linearised with the indicated enzymes, methylated with *Sss*I and religated. The purified circular DNA was then transfected into LMH/2A cells. In these substrates, the cleaved restriction site remained unmethylated after the circularisation. The ratio between methylated and unmethylated is shown. **F**, Luciferase assay using unmethylated VTG-CpGL in LMH/2A cells in which USF1 was depleted with siRNA; siLuc was used as control [NB: this siRNA does not recognise either of the luciferases expressed from our vectors]. Relative luciferase units (RLU) are defined as ratio between firefly and *Renilla* signal. The graphs show the mean ± SD of three independent experiments. **G**, RT-qPCR of *VTG* mRNA isolated from cells treated with siLuc or siUSF1 for 96 hours and EtOH or E_2_ for the last 24 h. The graph shows the mean ± SD of three independent experiments. Significance was assessed using Sidak’s multiple comparisons test. *P≤0.05, **P≤0.01, ***P≤0.001, ****P≤0.0001.

Having shown that methylation of the ten CpGs in the *VTG* enhancer/promoter by *Sss*I substantially attenuated E_2_ inducibility of the reporter (Fig. 5A), but that methylation of the ERE was without effect (Fig. 5B), we wanted to learn which CpGs were responsible for the transcriptional silencing. We therefore converted CpGs 1-8 (Fig. 1A) to TpGs by site-directed mutagensis. We could show that the C to T transition mutations at CpGs 1-6 and 8 failed to affect the basal transcriptional activity of the reporter and reduce its inducibility by E_2_. Indeed, the inducibility was increased in some cases (Fig. S5B). In contrast, C to T mutations in CpGs **c** and **d** in the ERE (mut ERE) and CpG7 (C594T) attenuated the inducibility upon E_2_ treatment (Fig. S5B,C). To learn whether the mutation in CpG7 affected USF1/2 binding, we tested the mutant oligo in an EMSA assay and found that the affinity of the protein for the mutated sequence was intermediate between that of the wt unmethylated CpG7 duplex and the methylated one (Fig. S5D). This translated directly to luciferase expression, where the mutant CpG7 (C594T) showed intermediate expression between the wt mock-methylated (wt C) and *Sss*I-methylated (wt mC) reporters (Fig. S5B). The expression of all the mutant reporters was inhibited upon methylation with *Sss*I, showing that the individual sites had no major influence on transcription of the reporter, irrespective of whether they were methylated or unmethylated (Fig. S5E).

In order to exclude the role of CpG1-6 and 8 in the transcriptional regulation of the VTG enhancer/promoter, we mutated them sequentially, such that the final mutant contained only CpG **c** and **d** in the ERE and CpG7. We tested the single and multiple mutants to exclude the possibility that interactions between different sites had additional influence on transcription, but this was not the case. All combinations tested were active when unmethylated and inhibited when methylated with *Sss*I (data not shown). Analysis of the final plasmid (ERE/7 wt) unmethylated and methylated confirmed that the silencing of the *VTG* enhancer/promoter was mediated by methylation of CpG7 in combination with methylation of the ERE (Fig. 5C), even though methylation of ERE alone had no effect on reporter activity (Fig. 5B).

In order to demonstrate the key importance of CpG7 in the control of *VTG* transcription, we methylated the wt reporter with *Eco72I* methylase (Rimseliene et al., 1995), which modifies solely the CACGTG E-box sequence. As shown in Fig. 5D, methylation of this site substantially attenuated basal luciferase expression, as well as its inducibility by E_2_, if not to the same extent as modification of all ten CpG sites by *Sss*I methylase. In contrast, methylation of the reporter with *Hpa*II methylase, which modifies CpG **d** in the ERE, had no appreciable effect on luciferase expression.

In a converse experiment, we wanted to reproduce the methylation pattern seen in the chicken embryo and in LMH/2A cells. To this end, we cleaved the reporter vector with *Pml*I, which cuts the E-box sequence and, as control, with *Hpa*II that cuts the ERE or *Xmn*I that cuts the vector backbone. We then methylated the linear DNA with *Sss*I and recircularised it with T4 DNA ligase. In the *Pml*I-cut vector, all CpGs with the exception of CpG7 were methylated, whereas only the *Hpa*II site in the ERE remained unmethylated in the *Hpa*II-cut vector and all CpGs were methylated in the *Xmn*I-cut vector. Transfection into LMH/2A cells showed that expression of the reporter luciferase gene could be efficiently induced by E_2_ solely when CpG7 was unmethylated (Fig. 5E, *Pml*I). In contrast, the reporter gene in the *Hpa*II-or *Xmn*I-cleaved vectors, as well as a plasmid that was treated with *Sss*I without prior cleavage (undigested) were refractory to induction with E_2_.

Finally, we wanted to confirm that the transcriptional control of *VTG* expression rests not only with CpG7, but also with the protein that binds the E-box: USF1/2. We therefore knocked down USF1 or luciferase (control) with siRNA (Fig. S5F-H) and transfected the cells with the reporter plasmids 48 hours later. Simultaneously, we assessed the induction of endogenous *VTG* in the siRNA-treated cells. Knock-down of USF1 reduced luciferase inducibility and the induction of endogenous *VTG* by more than 50% (Fig. 5F, G) or by a third in the case of luciferase inducibility with the second siRNA (Fig. S5H, left panel), which shows that USF1/2 plays a key role in the control of *VTG* transcription. (The incompleteness of the silencing could be explained by residual levels of the factor in the siRNA-treated cells.) This prompted us to conclude that USF1/2 binding to CpG7 is key to the regulation of expression of *VTG* and that its likely role in the prevention of methylation of its binding site is essential to ensure that the gene remains poised for ready hormone activation.

## DISCUSSION

We could confirm the findings of earlier studies and show that the chicken *VTG* gene was heavily-methylated already in early embryos, as well as in LMH/2A cells, and that it could be induced by a single dose of β-estradiol. The CpGs in the *VTG* enhancer/promoter seen to be demethylated in hormone-treated rooster liver (Saluz et al., 1986; Wilks et al., 1984; Wilks et al., 1982) remained resistant to bisulphite conversion in our experimental systems, however, we were able to detect low levels of the TET oxidation product hmC already 6 hours after E_2_ treatment that increased further with longer exposure to E_2_. The TET enzymes and TDG are present in both chicken embryonic and adult liver, as well as in the LMH2A cell line (data not shown), and it is therefore possible that the hormone-activated *VTG* gene is indeed targeted for demethylation, but that, unlike in the rooster, the demethylation machinery in our experimental set-up is not fully-functional. The finding that hmC proportion increased at the four CpGs upon hormone treatment at longer time points and that *VTG* induction was higher upon TET2 overexpression (data not shown) implied that the demethylation machinery can act on the *VTG* locus and that it has a positive effect on transcription. This could be further confirmed in cells treated with 5-azadeoxycytidine prior to induction with E_2_. In these cells, *VTG* expression was several-fold higher than even in the TET2-overexpressing cells (data not shown).

Puzzlingly, we failed to find evidence of hmC oxidation to fC or caC, steps necessary for the TDG-initiated base excision repair process that completes the multistep demethylation process. It is conceivable that the TET-mediated cascade terminated in our system after a single oxidation step, but this clearly wasn’t the case in the rooster experiments, where the genomic DNA became susceptible to cleavage with *Hpa*II that does not cleave hydroxymethylated DNA (Wilks et al., 1984; Wilks et al., 1982). Another possibility is that fC and caC excision and the subsequent BER steps were extremely rapid and were followed by immediate remethylation. Although such cyclic DNA demethylation/methylation events have been described in the pS2 promoter upon hormone treatment, it is unlikely that this was so in our system, because the cycling process supresses transcription (Kangaspeska et al., 2008; Metivier et al., 2008).

Although the above phenomenon requires further study, our data demonstrate that while global DNA demethylation facilitates transcription of the endogenous *VTG* gene upon hormone activation, site-specific demethylation of the ERE region is clearly not necessary. This agrees with our *in vitro* data (Fig. 2B) showing that the ERα binding to the ERE was unaffected by methylation. Indeed, the hormone-activated receptor needs to be able to bind to its cognate sequence irrespective of its methylation status, in order to recruit the chromatin-remodelling- and possibly also the demethylation machinery that are needed to facilitate transcription.

Given the requirement for ERα binding in E_2_-dependent *VTG* activation, the weak inducibility of the ΔG ERE reporter was puzzling, because the ΔG oligonucleotide did not detectably bind the receptor *in vitro* (Fig. 2B, C). However, it is possible that the observed induction was mediated by a second, imperfect ERE sequence, which lies around position −350 from the transcription start site (Fig. 1A).

As mentioned in the Introduction, the *VTG* enhancer/promoter has been studied extensively in the past and it could be shown that it contains several TF binding sites that control its basal- and liver-specific transcription, as well as its hormone-inducibility. Linker-scanning mutagenesis (Seal et al., 1991) revealed that deletion of the sequence between −113 and −335 eliminated the hepatocyte specificity, permitting the reporter contruct to be expressed also in fibroblasts. Importantly, insertion of a linker around position −50 abolished estrogen inducibility of the reporter. This mutation disrupted the E-box sequence, which contains CpG7 identified in this study as being critical for hormone induction of the silenced gene. Clearly, the interaction of the E-box binding factor (in our case the USF1/2 heterodimer) with the estrogen receptor is essential for hormone activation, given that mutational inactivation (Seal et al., 1991) or methylation (this study) of the E-box attenuates USF1/2 binding *in vitro* and the inducibility of the gene *in vivo*, as does mutation (but not methylation) of the ERE (Fig. S5B, C).

The interaction of different TF combinations bound at promoters and enhancers potentially provides cells with an enormous flexibility of gene expression, and DNA methylation extends this range still further by altering the affinity of some, but not all, TFs for their respective recognition sites. Moreover, the E-box hexamer consensus sequence CANNTG has been shown to bind a large family of basic helix-loop-helix leucine zipper (b-HLH-LZ) proteins that can bind as homo-or heterodimers, and often in methylation-sensitive manner (Fujii et al., 2006; Hou et al., 2012; Perini et al., 2005; Prendergast et al., 1991; Prendergast and Ziff, 1991). Yet, in spite of the multitude of proteins that could potentially bind to the *VTG* E-box (see Table 1) and the plethora of possible protein/protein interactions (both activating and repressing) that this could generate, *VTG* expression is strictly-controlled and appears to require exclusively USF1/2. This was unexpected. USF1/2 has been reported to interact with ERα and to bind preferentially to the perfect E-box sequence CACGTG and less so to CATGTG (Corre and Galibert, 2005; deGraffenried et al., 2004). This could explain why the C to T mutation in CpG7 had such a deleterious effect on estrogen inducibility (Fig. S5B, C). However, it does not explain why the factor bound so efficiently to the different sequences in our EMSA assays (Fig. 2D,E; Fig. S2F,G,H). Similarly, methylation of the E-box sequence would also lower the E_2_ inducibility of the VTG enhancer/promoter, because the affinity of USF1/2 for the methylated sequence is low (Fig. 2D, E). As noted above, however, there are a number of TFs that could potentially bind to the E-box in the absence of USF1/2 and activate VTG transcription, particularly as some – notably c-Myc – have been reported to interact with ERα (Cheng et al., 2006). These questions require clarification in the future, as does the reason why the *VTG* E-box sequence is the only CpG site out of the ten that remains unmethylated *in vivo*. It is possible that its methylation is actively prevented, possibly due to the fact that it is rapidly bound by USF1/2 or other E-box proteins and thus that the access of methyl transferases to this site is hindered. This has been proposed as a possible mechanism that protects CpG islands from *de novo* methylation (Brandeis et al., 1994; Macleod et al., 1994), but whether inducible enhancers/promoters where only a single site is affected are subject to similar protection from methylation remains to be demonstrated.

DNA methylation changes, particularly hypermethylation, have been associated with a long list of pathologies and disorders, ranging from cancer, immune system dysegulation (Crohn’s disease, Grave’s disease, rheumatoid arthritis, type 1 diabetes), neurological disorders (Alzheimer’s disease, Fragile X syndrome, schizophrenia, epilepsy, depression) and atherosclerosis to osteoporosis (Ehrlich, 2019). In the vast majority of the studies concerned with the human methylome, focus was placed on CpG islands and other CpG-rich regions, the hypermethylation (or indeed hypomethylation) could be seen to be associated with the given pathology. Our study highlights an important caveat of these studies, namely, that the alteration of the methylation status of a single CpG can have equally dramatic consequences as the hypermethylation of a CpG island. The outcome of both events can be the silencing – or at least attenuation - of transcription. Interestingly, the two main protagonists of this study, ERα and USF1/2, may be implicated (separately and together) in a subset of the above syndromes. USF1/2 plays a major role in cell proliferation control, as well as in the control of expression of the *TP53* (Hale and Braithwaite, 1995; Reisman and Rotter, 1993) and *BRCA2* genes but it has been reported to be transcriptionally inactive in three of six transformed breast cancer cell lines (Ismail et al., 1999), the widely-used MCF7 line among them, which has a hypermethylator phenotype (Xing and Archer, 1998). The two TFs interact in the transcriptional activation of the Cathepsin D gene, the dysregulation of which is linked to a number of the above-listed disorders (Dubey and Luqman, 2017). It will be interesting to learn whether some of these dysregulation events might be linked to the aberrant methylation of USF1/2/ E-boxes within the transcriptional control regions of genes involved in these pathologies.

## MATERIALS AND METHODS

### Cell culture

LMH/2A cells (ATCC CRL-2118) were grown in Williams’ E medium (GIBCO) without phenol red, supplemented with 10% charcoal-stripped FCS, streptomycin/penicillin (100 U/ml) and L-glutamine (2.4 mM). MCF-7 cells were grown in DMEM (GIBCO) with phenol red, supplemented with 10% FCS, streptomycin/penicillin (100 U/ml) and insulin (Sigma-Aldrich, 10 µg/ml). Cell lines were grown at 37°C in a 6% CO_2_ humidified atmosphere. Where indicated, cells were treated with 100 nM β-estradiol (E_2_, ethanol solution, Sigma-Aldrich) or an equivalent volume of ethanol, or with 8 µM aphidicolin (aph, DMSO solution, Sigma-Aldrich) or an equivalent volume of DMSO. E_2_ and/or aphidicolin were added fresh every 12 h, because of instability under cell culture conditions. RNA interference was carried out using Lipofectamine RNAiMAX (ThermoFisher) according to the manufacturer’s instructions. SiRNA against chicken 1-USF1: ^5’^CCCAAUAUCAAAUAUGUCUUC^3’^; 2-USF1: ^5’^UAUGUCUUCCGCACAGAGAUU ^3’^; against luciferase (control): ^5’^CGUACGCGGAAUACUUCGAdTdT^3’^.

### Handling and treatment of fertilized eggs

Eggs were prepared as described (Baeriswyl and Stoeckli, 2006) and kept at 39°C. At day 9, the eggs were treated with ethanol or 100 nM E_2_ by pipetting ∼55 µl (depending on weight of egg) of a 100 µM solution (10% EtOH) into the egg, through the window prepared at day 3 of development. 24 hours later, the embryos were sacrificed, the livers were excised and immersed in RNA*later* (Sigma). Livers were homogenized using a Tissue Ruptor (Qiagen) in RIPA buffer for proteins and in RLT buffer for RNA. Following homogenization, protein samples were processed further according to the Western Blot protocol and RNA was extracted using an RNAeasy Kit (Qiagen).

### Western Blotting (WB)

Cells were collected using trypsin, washed in phosphate-buffered saline (PBS) and lysed in RIPA buffer (50 mM Tris pH 8, 1 mM EDTA, 1% NP-40, 0.5% deoxycholate (DOC), 0.1% SDS, 150 mM NaCl). Lysates were sonicated (18 s, 50 cycles, 70% amplitude; in a Bandelin Sonoplus GM70) and protein concentrations were measured using the Bradford assay. 30-80 µg of protein were boiled in 1x SDS loading buffer (50 mM Tris, pH 6.8, 10% glycerol, 1.6% SDS, 0.1 M DTT, 0.01% bromophenol blue) and separated on polyacrylamide gels (6-10%) in 10% SDS-running buffer at 130 V. Proteins were transferred onto activated PVDF membranes (Amersham Pharmacia Biotech) which were incubated with primary antibody overnight at 4°C in 5% milk, washed 3x with TBS-T and incubated for 1 hours at RT with the secondary antibody (horseradish peroxidase-conjugated sheep anti-mouse or donkey anti-rabbit IgG, GE Healthcare). Membranes were analyzed with a Fusion Solo (Vilber Lourmat). Antibodies used were mouse Anti-VTG (Abcam, ab36794), rabbit Anti-TFIIH p89 (Santa Cruz, sc-293), mouse Anti-ERα (ThermoFisher, MA5-13065), rabbit Anti-USF1 (GeneTex GTX16396), mouse Anti-USF2 (Santa Cruz, sc-293443).

### Real-time quantitative PCR (RT-qPCR)

Cells were harvested using trypsin and RNA was extracted according to manufacturer’s instructions (RNeasy, Qiagen). 2 µg of the extracted RNA were reverse-transcribed using the high capacity cDNA reverse transcription kit (Applied Biosystems) according to manufacturer’s instructions. 125 ng of the cDNA were used for the PCR using the platinum SYBR green qPCR superMix-UDG kit (Invitrogen) according to manufacturer’s instructions, except that the reaction was scaled down from 50 µl to 20 µl. The standard cycling program for ABI instruments was used (50°C for 2 min, 95°C for 2 min, 40x (95°C for 15 s, 60°C for 30 s) and melting curve analysis at 65°C for 15s, heating to 97°C with continuous acquisition per 5°C, 40°C for 30 s). The RT-qPCR was run on a LightCycler 480 (Roche). GAPDH primers were used as an internal control. Technical triplicates were made for every sample and primer. The primers used are listed in Table S1 (SI).

### Chromatin immunoprecipitation

75 Mio LMH/2A cells were harvested after 24 hours treatment with 100 nM E_2_ or EtOH. Cells were washed twice with PBS and fixed for 10 min in 1% formaldehyde at RT. Fixation was quenched with 125 mM glycine at RT for 10 min. Cells were collected by centrifugation at 300 x g for 5 min at 4°C, washed twice with PBS and the pellet was resuspended in cell lysis buffer (10 mM Tris, pH 8, 1 mM EDTA, 0.5% IGEPAL, protease inhibitors) and incubated on ice for 10 min. The lysate was centrifuged at 2’000 x g at 4°C for 5 min. The pellet was resuspended in nuclear lysis buffer (10 mM Tris, pH 8, 1 mM EDTA, 0.5 M NaCl, 1% Triton X-100, 0.5% sodium deoxycholate, 0.5% lauroylsarcosine, protease inhibitors) and incubated on ice for 10 min with repeated vortexing. Lysate was centrifuged at 3’000 x g for 5 min at 4°C. The pellet was resuspended in PBS, split for different immunoprecipitations and centrifuged at 3’000 x g for 10 min at 4°C. Pellets were resuspended in 300 µl lysis buffer (10 mM Tris, pH 8, 1 mM EDTA, 150 mM NaCl, 0.1% sodium deoxycholate, 0.1% SDS, protease inhibitors) and sonicated at maximum power for 10 min with 30 s intervals in a Bioruptor (Diagnode). Samples were centrifuged at 16’000 x g for 10 min at 4°C and the supernatant was collected. Input samples (15%) were frozen and the rest was made up to 1 ml with IP buffer (16.7 mM Tris, pH 8, 1.2 mM EDTA, 300 mM NaCl, 1.1% Triton X-100, protease inhibitors). 5 µg of the respective antibodies (Anti-ERα; ThermoFisher MA5-13065 or Anti-Flag; Sigma F3165) were added to the lysate and incubated on a rotating wheel at 4°C overnight.

The beads (G protein Sepharose, GE Healthcare) were washed twice in IP buffer and centrifuged for 2 min at 2’700 x g before blocking with 1 mg/ml BSA in IP buffer for 1 h. After one additional wash, 50 µl of beads were added to 1 ml lysate with the antibody. The mixture was incubated for 3 hours on a rotating wheel at 4°C. Mixtures were centrifuged for 2 min at 2’000 x g and the supernatant was removed. The beads were washed successively in 1 ml of four different wash buffers for 5 min on the rotating wheel (wash buffer 1: 20 mM Tris, pH 8, 2 mM EDTA, 0.1% SDS, 1% Triton X-100, 300 mM NaCl; wash buffer 2: like wash buffer 1, but with 500 mM NaCl; wash buffer 3: 10 mM Tris, pH 8, 1 mM EDTA, 1% sodium deoxycholate, 500 mM LiCl, 1% NP-40; wash buffer 4: 10 mM Tris, pH 8, 1 mM EDTA). One additional wash with wash buffer 4 was performed, during which the beads were divided for DNA and proteins. For proteins: the beads were taken up in 2x SDS loading dye (100 mM Tris, pH 6.8, 20% glycerol, 3.2% SDS, 0.2 mM DTT, 0.04% bromophenol blue), the mixture was vortexed vigorously and incubated for 30 min at 100°C (denaturing and de-cross-linking). The beads were spun down and the supernatant was loaded on an SDS-PAGE together with inputs. Gels were processed further according to general WB protocol. For DNA: beads and inputs were taken up in 100 µl crosslink reversal buffer (20 mM Tris, pH 8, 0.5 mM EDTA, 0.1 M NaHCO_3_, 1% SDS, RNase at 10 µg/ml (Roche)) and incubated at 65°C overnight. The next day, the DNA was purified using the PCR purification kit from Qiagen and eluted in 10 µl of H_2_O. RT-PCR was performed on 3.5 µl of DNA using the platinum SYBR green qPCR superMix-UDG kit (Invitrogen) according to manufacturer’s instructions, except that the reaction was scaled down to 20 µl and 1 mM of MgCl_2_ were added. The following protocol was used for amplifications: 50°C for 2 min, 95°C for 2 min, 50x (95°C for 15 s, 57°C for 30 s) and melting curve analysis at 65°C for 15 s, heating to 97°C with continuous acquisition per 5°C, 40°C for 30 s). The qPCR was run on a LightCycler 480 (Roche) with the ChIP primers listed in table S1.

### Labelling of oligos

Fill-in reaction (3’-labelling): 2 pmoles of oligo duplex with a 5’-overhang were incubated for 15 min at RT with 0.2 µl of the respective [α-^32^P]-dNTP (2 µCi, Hartman Analytic), 1 mM of each of the other three dNTPs and 5 U Klenow fragment (3’—>5’ exo-, NEB) in the corresponding buffer. The enzyme was heat-inactivated for 20 min at 75°C and the reaction was allowed to slowly cool down to re-anneal the strands. The free dNTPs were removed on a Sephadex G-25 column (GE Healthcare).

Kinase reaction (5’-labelling): 2 pmoles of ssDNA were phosphorylated with 0.7 µl of [γ-^32^P]-ATP (7 µCi, Hartman Analytic) using T4 polynucleotide kinase (10 U, NEB). The reaction was supplemented with 10 mM DTT and incubated for 30 min at 37°C. The reaction was heat-inactivated for 5 min at 95°C and the free nucleotides were removed on a Sephadex G-25 column (GE Healthcare). The oligo was annealed to a 1.5x excess of the unlabelled complementary strand by heating for 5 min to 80°C and slowly cooling to RT in annealing buffer (50 mM Hepes, pH 7.5, 100 mM NaCl).

### Nuclear extracts

All nuclear extracts were prepared according to Dignam et al. (Dignam et al., 1983), with or without pre-treatment of the cells with E_2_.

### Electrophoretic Mobility Shift Assay (EMSA)

All EMSA reactions were carried out in binding buffer (10 mM Tris-HCl, pH 7.6, 50 mM NaCl, 1 mM DTT, 1 mg/ml BSA, 5% glycerol (Hyder et al., 1999)) in a volume of 5 µl using the unspecific competitor poly(dI-dC) (Sigma-Aldrich). 20 ng of poly(dI-dC) were used for 100 ng of recombinant ERα (ThermoFisher, RP-310) and 1 µg for 10 µg of nuclear extracts. For all EMSAs, 10 fmol of [α-^32^P]-dNTP labelled, or [γ-^32^P]-ATP phosphorylated oligos were used. Where indicated, an excess of unlabelled competitor DNA or 100 nM of E_2_ were added. For the supershift assays, 200 ng of antibody (Anti-ERα; ThermoFisher MA5-13065) were added together with the proteins. Proteins, and competitor or antibodies where indicated, were incubated for 20 min on ice in binding buffer. Labelled DNA was added and the mixture was left at RT for another 20 min before loading on a 5% polyacrylamide gel (Acryl/ Bis 29:1, Amresco) eluted with 1x TAE (40 mM Tris, 20 mM acetate, 1 mM EDTA). The gels were run at 200 V for 1 hours in 1x TAE and dried in a gel dryer for 1 h. The gels were exposed to phosphor screens and the autoradiographs were developed in a Typhoon FLA 9500 (GE Healthcare Life Sciences). The oligonucleotides used are listed in Table S2 (SI).

### SDS-PAGE of cross-linked binding reactions

EMSAs were performed as described and cross-linked for 5 min (∼720 mJ) in a UV Stratalinker 1800 (Stratagene). Samples were taken up in 2x SDS loading buffer and boiled for 5 min before being loaded on a 7.5 % denaturing polyacrylamide gel and run for 1-2 hours in 10 % SDS-running buffer at 130 V. Gels were dried for 1 h, exposed to phosphor screens and developed in a Typhoon FLA 9500 (GE Healthcare).

### CpGL cloning and Luciferase assay

The enhancer/promoter region of the chicken vitellogenin II gene spanning nucleotides −638 to −1 was cloned into a CpG-free firefly luciferase plasmid (pCpGL-basic, InvivoGen) using *Hin*dIII and *Nco*I. The enhancer/promoter region was amplified from genomic DNA of LMH/2A cells using the primers ‘enhancer fwd and rev’ (Table S1, SI). The plasmid was further cut with *Hin*dIII and *Hpa*II to excise the intervening sequence and the annealed upper

^5^’AGCTTAAAAATATTCCTGGTCAGCGTGAC3’ and lower

^5’^CGGTCACGCTGACCAGGAATATTTTTA^3’^ ds-oligo was ligated into the plasmid to translocate the *Hin*dIII site closer to the ERE for further ligations of the ERE. On the other side of the ERE a *Kpn*I site was created by mutagenesis and another *Kpn*I site in the plasmid had to be destroyed by mutagenesis (primers in Table 1, SI).

The resulting VTG-CpGL plasmid was co-transfected with a Renilla luciferase-expressing plasmid (pRL-SV40, Promega) into LMH/2A cells using Lipofectamine 2000 (Invitrogen) according to manufacturer’s instructions. Cells were re-seeded the next day in technical triplicates both for E_2_ and ethanol treatment and treated for 24 h. Cells were subsequently lysed directly in firefly luciferase substrate from Promega (Dual-Glo Luciferase kit) for 30 min, with vigorous shaking.

Cell lysates were processed further according to manufacturer’s instructions and luminescence was measured on the SpectraMax i3 (Molecular Devices) plate reader. Relative luciferase units (RLU) were calculated as the ratio between the firefly and *Renilla* signals.

### Mutagenesis of VTG-CpGL

200 ng of vector DNA were mixed with 50 pmoles forward and reverse primers, 250 µM dNTPs, 2% DMSO and 1 U Phusion high fidelty polymerase (NEB) in 1x buffer provided by the manufacturer. Extension was performed in 50 µl according to the following protocol: 95 °C, 2 min; 30x (95 °C, 1 min, 55 °C, 1 min, 63 °C, 30 min); 68 °C, 20 min; 15 °C hold. Extension reaction was digested twice with 20 U *Dpn*I for 1 hours at 37°C followed by heat inactivation of the enzyme for 20 min at 80°C. The reaction mix was then transformed into electro-competent Pir1 bacteria (R6K gamma ORI, Invitrogen) after being desalted by incubating for 15 min on a 0.025 µm Millipore nitrocellulose filter on H_2_O. Bacteria were shaken at 37°C for 40 min and plated onto Zeocin (25 µg/ml, InvivoGen) agar plates. Clones were picked the next day, grown in liquid culture overnight and plasmids were extracted using the NucleoSpin Plasmid Kit (Macherey-Nagel) according to manufacturer’s instructions. Plasmids were sequenced at Microsynth to check for successful mutagenesis. The primers for mutagenesis are listed in Table S1 (SI).

### Ligation of modified ERE into VTG-CpGL

50 µg of the plasmid DNA were cut with *Hin*dIII-HF (NEB) in Cut Smart buffer with 100 U of enzyme for 1.5 hours at 37°C in 50 µl. Linearization was verified on a 1% agarose gel and enzyme was heat-inactivated for 20 min at 80°C. For the first ligation step, 10 µg of linearized DNA and 20x excess of annealed oligo with different modifications at the CpGs (ERE insert, Table S1, SI) were incubated for 2 hours at RT in T4 ligase buffer with 400 U of T4 ligase (NEB) in a total volume of 100 µl. Ligase was inactivated by heating to 65°C for 10 min and efficient ligation was verified on an agarose gel. Reaction volume was increased to 200 µl with 1x Cut Smart buffer, 40 U of *Kpn*I-HF (NEB) were added and incubation was continued for additional 1.5 hours at 37°C. 20 U of *Hin*dIII-HF were then added, the mixture was incubated for an additional hour and subsequently cleaned-up on MinElute columns (5µg per column, Qiagen) and eluted in 20 µl H_2_O. 20 reactions of the first ligation were pooled for the second ligation. 12.5 µg of DNA from the first ligation were incubated in 10 ml of 1x T4 ligase buffer with 4000 U of ligase for 1 hours at RT. Recircularization was verified on an agarose gel. The resulting mix was concentrated by EtOH precipitation and the supercoiled form of the plasmid was extracted on a CsCl gradient and purified further as described (Baerenfaller et al., 2006).

### Expression and purification of Eco72IM

The *Eco*72IM sequence was PCR-amplified from the *Eco*72IRM plasmid (Thermo Fisher) and recombined with the pDONR 221 (Gateway, Thermo Fisher) to form an entry vector. The resulting entry vector was recombined with pDest15 (Gateway) adding an N-term GST tag to the methyltransferase. BL21 pLysS *E. coli* (Promega) were transformed with the expression vector. A starter culture (5 ml) was grown from a single colony and subsequently used to inoculate a 1 l culture that was grown at 37°C to an OD of 0.65. Culture was cooled down to 22°C, a sample of uninduced culture was taken for SDS gel and protein expression was induced with the addition of 250 µM IPTG with shaking overnight. The next day, bacteria were washed with cold PBS by centrifugation at 4’000 x g for 10 min at 4 °C. 14 ml lysis buffer (100 μg/ml Lysozyme, 1 % Triton, 1 mM PMSF, 1 x protease inhibitor (cOmplete, Roche), 10 mM DT, 1 x PBS) were added to pellet and lysate was incubated for 30 min in a beaker stirring at 4 °C. Lysate was sonicated twice for 1 min on ice (ampl 70%, 50% cycle), together with uninduced sample for SDS gel and centrifuged at 18’000 x g for 30 min at 4 °C. GSH beads (GE Healthcare, 600 µl) washed in lysis buffer were added to the supernatant and incubated on a rotating wheel for 2 hours at 4 °C. The beads were then washed three times in washing buffer (10% glycerol, 10 mM DTT, 1 mM PMSF, 1 x PBS) by inverting the tube several times followed by centrifugation at 500 x g for 5 min. Flow-through and first wash were retained for the SDS gel. The beads were then incubated with 500 µl elution buffer (washing buffer with freshly-added 20 mM glutathione, pH adjusted to 8 with NaOH) on rotating wheel at 4 °C. The first elution was aliquoted and snap frozen.

### In vitro methylation

1 µg of plasmid DNA was incubated in 1x NEB buffer 2 (*SssI*) or Eco72IM buffer (10 mM Tris HCl, 50 mM NaCl, 1 mM DTT, 10 mM EDTA), 160 µM freshly-diluted SAM (NEB) and 4 U of *Sss*I (NEB) or 4 µl purified *Eco*72IM (diluted 1:100) for 1 hours at 37°C. The reaction was stopped by heating to 65°C for 20 min.

### EdU incorporation and Click-iT Reaction

The cells were grown on coverslips and treated with 8 µM aphidicolin or DMSO, 100 nM E_2_ or ethanol and 10 µM of 5-ethynyl-2’-deoxyuridine (EdU, Invitrogen) for 24h. Medium was removed, the cells were washed once with PBS and 1 ml of 3.7% formaldehyde in PBS was added. The cells were fixed for 15 min at RT, then washed twice with 1 ml of 3% BSA in PBS. 1 ml of 0.5% Triton X-100 in PBS was added and the cells were incubated for 20 min at RT. Click-iT reaction master mix was prepared. [For 5 slides: 129 µl 1x Click-iT reaction buffer (freshly diluted 1:10 in H_2_O), 6 µl CuSO_4_, 0.36 µl AlexaFluor azide, 15 µl reaction buffer additive (freshly diluted 1:10 in H_2_O)]. Permeabilization buffer was removed and the cells were washed twice with 1 ml 3% BSA in PBS. 30 µl of Click-iT reaction mix were pipetted onto Parafilm and the coverslips were put cells-down onto the mix and incubated for 30 min at RT, protected from light. The coverslips were washed once with 1 ml of 3% BSA in PBS and then washed well with 1x PBS to remove BSA. After one final wash with H_2_O, the cells were fixed with mounting media containing 4’,6-diamidino-2-phenylindole (DAPI, VectaShield). The slides were analyzed on an Olympus IX81 fluorescence microscope.

### DNA extraction, oxidation of hmC and bisulphite conversion

Genomic DNA was extracted from cells using the Wizard Genomic DNA Purification Kit (Promega). DNA was eluted with water and digested with *Eco*RV overnight. EtOH precipitation was performed and DNA was additionally cleaned up on Micro Bio-Spin 6 chromatography columns in SSC (BioRad) as purity was essential for oxidation of hmC. Samples were split and one half was subjected to oxidation. Selective oxidation of hmC to fC was achieved using potassium perruthenate (KRuO_4_, Sigma-Aldrich) as described (Booth et al., 2012; Booth et al., 2013). In short, 0.5-2 µg of DNA were incubated in 50 mM NaOH in 24 µl for 30 min at 37°C after vigorous vortexing to denature DNA. 1 µl of 15 mM KRuO_4_ solution in 50 mM NaOH was added and oxidation was incubated on ice for 1 hours with vortexing every 5 min. Reaction was cleaned up on polyacrylamide columns (89849, ThermoScientific) and processed further with the rest of the sample for bisulphite conversion. Bisulphite conversion was achieved using the EZ DNA Methylation-Gold Kit from Zymo Research according to manufacturer’s instructions. The cycling protocol was adapted because of the slightly less efficient conversion of fC as compared to unmodified cytosine (95°C for 5 min; 2x (60°C for 25 min, 95°C for 5 min, 60°C for 85 min, 95°C for 5 min, 60°C for 175 min, 95°C for 5 min); 20°C hold.

### Polymerase chain reaction on bisulphite-converted DNA

100 ng of bisulphite-converted DNA was used as template for the PCR using the ZymoTaq DNA polymerase (Zymo Research). The reaction was carried out in a total volume of 40 µl in the reaction buffer provided by the manufacturer supplemented with of 1 mM dNTPs, 1.5 mM MgCl_2_, 500 mM each primer and 2 U of polymerase. The cycling protocol was as follows: 10 min 95°C; 5x (30 s 94°C; 30 s 52°C; 90 s 72°C); 5x (30 s 94°C; 30 s 52°C; 90 s 72°C); 35x (30 s 94°C; 30 s 55°C; 90 s 72°C); 7 min 72°C. The primers used for the amplification of bisulphite converted DNA are listed in Table S1 (SI). The forward primers were only added after the first 5 cycles to reduce the formation of primer dimers. The PCR fragments were purified using a BluePippin (Sage Science) on a 2% agarose gel according to manufacturer’s instructions, before being bar-coded for PacBio sequencing.

### PacBio sequencing and analysis

The PacBio single-molecule real-time (SMRT) sequencing technology works by a strand-displacement mechanism on circularised single molecules. The highly-processive polymerase copies the circle multiple times to generate a long read containing many repeats of the same sequence. The deconvolution to single reads generates a consensus sequence that corrects the high error-rate of the polymerase, resulting thus in very accurate sequencing results. The sequencing and data evaluation were carried out in collaboration with the Functional Genomic Center Zurich (www.fgcz.ch).

### Affinity purification and mass spectrometry

Different CpG-containing oligonucleotides with 5’-TTAA overhangs were annealed and end-to-end ligated overnight at 16°C with T4 ligase (NEB). Oligos were ethanol-precipitated and filled-in with 1 mM biotinylated dUTP (Thermo Fisher) using Klenow fragment (NEB). Reactions were passed twice through a Sephadex G-25 desalting column (GE Healthcare) to get rid to free dUTP and subsequently bound to Dynabeads M-280 Streptavidin (Invitrogen) by rotating 20 min at RT. 1.5 mg of nuclear extracts were pre-incubated with 30 µg of poly(dI-dC), before an equal volume was added to the beads. Binding was allowed to take place in 1x EMSA buffer without BSA for 30 min on a rotating wheel at 4 °C. Beads were washed 2x with 60 µl 1x EMSA buffer without BSA. Bound proteins were eluted with 60 µl of elution buffer (10 mM Tris-HCl pH 7.6, 5 % glycerol, 1M NaCl) 30 min rotating at 4°C. Elution was desalted on 0.025 µm VSWP membranes (Merck) against water for EMSAs.

### Shotgun LC-MS/MS

The protein mixture was digested with trypsin. After cleaning up, the peptide mixture was applied to a reversed phase high-performance liquid chromatography (HPLC) column and separated prior to ionization and analysis by the mass spectrometer. The peptide ions selected by an instrument algorithm for fragmentation were recorded as a peptide signature, which was analyzed by a sequence algorithm (MASCOT). The peptides were scored according to the average probability using total spectra counts. The complete list of the MS results can be found online in Supplementary Information.

### Immunodepletion of extracts

50µg Dynabeads Protein G (Invitrogen), washed in PBS-T (0.1% Tween 20), were incubated with 0.4 µg of the respective antibody for 20 min at RT on rotation wheel. Beads were washed twice in PBS-T by inverting the tube several times and putting it on the magnet for 2 min. 100 µg of nuclear extracts were added and binding was allowed for 2 hours rotating at 4 °C. Supernatant was aliquoted and snap frozen as immunodepleted extract. Samples were taken for control Western Blots and EMSAs.

### Statistical analysis

All experiments were performed at least three times. Results are shown as means +/-SD. Statistical significance was determined byStudent’s *t*-tests. P ≤ 0.05 was considered statistically significant.

## ACKNOWLEDGEMENTS

The authors would like to express their gratitude to Hanspeter Saluz for providing hen and rooster DNA samples, to Peter Hunziker for the proteomic analysis and Giancarlo Russo for assistance with the bioinformatics, to Thermo Fischer for the generous gift of the Eco72I DNA, to Beat Kunz for assistance with the egg manipulations and to Maite Olivera Harris for constructive discussions.

